# Spatial working memory representations in human cortex are robust to a task-irrelevant interrupting stimulus

**DOI:** 10.1101/2021.09.16.460692

**Authors:** Kelvin Vu-Cheung, Edward F Ester, Thomas C Sprague

## Abstract

Visual working memory (WM) enables the maintenance and manipulation of information no longer accessible in the visual world. Previous research has identified spatial WM representations in activation patterns in visual, parietal, and frontal cortex. In natural vision, the period between the encoding of information into WM and the time when it is used to guide behavior (the delay period) is rarely “empty”, as is the case in most of the above laboratory experiments. In naturalistic conditions, eye movements, movement of the individual, and events in the environment result in visual signals which may overwrite or impair the fidelity of WM representations, especially in early sensory cortices. Here, we evaluated the extent to which a brief, irrelevant interrupting visual stimulus presented during a spatial WM delay period impaired behavioral performance and WM representation fidelity assayed using an image reconstruction technique (inverted encoding model). On each trial, participants (both sexes) viewed two target dots and were immediately post-cued to remember the precise spatial position of one dot. On 50% of trials, a brief interrupter stimulus appeared. While we observed strong transient univariate visual responses to the distracter stimulus, we saw no change in reconstructed neural WM representations under distraction, nor a change in behavioral performance on a continuous recall task. This suggests that spatial WM representations may be particularly robust to interference from incoming task-irrelevant visual information, perhaps related to their role in guiding movements.

## Introduction

Working memory (WM) is the ability to store and manipulate information over brief periods to accomplish behavioral goals. Maintaining spatial locations in WM results in sustained spiking activity in macaque prefrontal cortex (PFC) (Funahashi et al., 1989; Fuster and Alexander, 1971; Kubota and Niki, 1971), parietal cortex (Chafee and Goldman-Rakic, 1998; Gnadt and Andersen, 1988), and visual cortex (Supèr et al., 2001; van Kerkoerle et al., 2017). Similar elevated delay-period activity in PFC, parietal, and visual cortex was also found using fMRI (Courtney et al., 1997; Curtis and D’Esposito, 2003; Leung et al., 2002; Saber et al., 2015; Srimal and Curtis, 2008; reviewed in Curtis and Sprague, 2021). Subsequent studies demonstrated that sustained activity patterns in these regions encode the content of working memory, as it was possible to decode remembered spatial positions (Hallenbeck et al., 2021; Henderson et al., 2021; Jerde et al., 2012; Li et al., 2021; Sprague et al., 2016, 2014), orientations (Christophel et al., 2018; Ester et al., 2015, 2013, 2009; Harrison and Tong, 2009; Rademaker et al., 2019; Serences et al., 2009; Yu and Shim, 2017), colors (Serences et al., 2009; Yu and Shim, 2017), motion directions (Christophel and Haynes, 2014; Emrich et al., 2013; LaRocque et al., 2017; Riggall and Postle, 2012), and complex patterns (Christophel et al., 2015, 2012) from activation patterns in prefrontal, parietal, and visual cortex (reviewed in Christophel et al., 2017; Serences, 2016). However, in many of these studies, the WM delay period remained blank, leaving open the possibility that visual stimulation could disrupt the sustained neural representation held in WM.

An alternative neural coding scheme has also been supported by recent computational modeling, EEG and MEG studies whereby WM representations are stored as brief changes in local synaptic efficacy (Mongillo et al., 2008; Rose et al., 2016; Wolff et al., 2020, 2017, 2015). This ‘activity silent’ coding scheme does not necessitate persistent spiking activity, and instead predicts that transient changes to the local neural network can be ‘read out’ using a barrage of neural input, or, in an experimental setting, using an irrelevant stimulus to ‘ping’ the state of the network (Stokes, 2015). In one example study, participants remembered the orientation of a grating, and on some trials a large high contrast task-irrelevant stimulus flashed briefly during the delay period. The impulse evoked a transient EEG activity pattern which contained a representation of the remembered orientation which was absent prior to the impulse (Wolff et al., 2015). In an fMRI experiment, which measures hemodynamic signals over a slower timescale, could an intervening stimulus lead to a similar evoked enhancement of WM representations? Or, does the intervening stimulus instead act to overwrite neural signals measured with fMRI, resulting in a disrupted neural code?

Behavioral and neural studies have demonstrated that WM is remarkably resistant to most interrupters (intervening stimuli presented during the maintenance period of WM tasks, as defined by Hakim et al., 2020). As an example, participants in an fMRI study performed a delayed orientation recall task while either low intensity interrupters (static grating or noise, which did not disrupt behavioral performance) or high intensity interrupters (flashing gratings, noise, faces or gazebos, which did disrupt performance) were presented throughout the delay. The ‘intensity’ of the behavioral distraction predicted disruptions in neural WM representations (Rademaker et al., 2019). There are several instances where neural working memory representations are impacted by an interrupter – when performance is impaired, WM representations follow (Rademaker et al., 2019; but see Bettencourt and Xu, 2016), and when performance is biased by a similar interrupter, so are neural representations in visual cortex (Hallenbeck et al., 2021; Lorenc et al., 2018). However, these studies have not tested whether a spatial WM stimulus is robust to a task-irrelevant brief interrupter, like that used to activate activity-silent WM representations in previous work. Such a stimulus is important to test because, if activity-silent representations play a meaningful role in supporting spatial WM, the interrupter may instead enhance observed neural WM representations (Stokes, 2015; Wolff et al., 2017, 2015).

Here, we tested the impact of a task-irrelevant interrupting stimulus on the quality of spatial WM representations. If spatial WM representations are resistant to disruption, we expect to see no change in WM representation quality following task-irrelevant interruption. However, if activity-silent codes contribute to spatial WM representations, we would expect to see a transient enhancement in their fidelity. We found that spatial WM representations were remarkably stable, with no evidence for either a transient disruption or augmentation due to an interrupting stimulus.

## Materials & Methods

### Participants

Human participants were recruited from the UC San Diego community and provided written consent. All procedures were approved by the UC San Diego Institutional Review Board. All procedures were approved by the [withheld] Institutional Review Board. N = 4 participants were recruited for the study (3 female; aged 25-30 yrs), and no participants were excluded. Participants received cash compensation ($20/hr fMRI scans, $10/hr behavioral testing). Sample size was not pre-determined and is similar to that used for similar studies (Sprague et al., 2014, n = 4).

### Experimental procedures: overview

We aimed to compare spatial WM representations carried by neural activation patterns in human retinotopic cortex between conditions requiring participants maintain a single spatial location in WM over a blank delay interval and over a delay interval with a brief task-irrelevant interruption. Participants performed one 1-hr behavioral session to familiarize themselves with the behavioral tasks performed inside the scanner, one 2-hr fMRI scanning session to acquire anatomical images and retinotopic mapping data, and two 2-hr fMRI scanning sessions to acquire data for the spatial WM task and spatial mapping task. During each fMRI session, participants performed 8 runs of the spatial WM task and 4 runs of the spatial mapping task. All fMRI data was analyzed within-session and combined across sessions and participants for analyses. We also acquired fMRI data while participants performed a general spatial localizer task used to select voxels for further analysis.

We presented stimuli using the Psychophysics toolbox (Brainard, 1997; Pelli, 1997) for MATLAB (The Mathworks, Natick, Mass). During behavioral practice sessions, participants viewed stimuli on a contrast-linearized LCD monitor (1920×1080, 60 Hz) positioned 62 cm away. Participants were comfortably seated in a dimmed room and positioned using a chin rest. During scanning, participants viewed all stimuli through a mirror mounted on the fMRI head coil. Stimuli were projected on a screen placed at the foot of the scanner bore (viewing distance: 370 cm; screen width: 120 cm) using a contrast-linearized LCD projector at a resolution of 1024 × 768 at 60 Hz refresh rate.

### Task design: spatial WM task

Throughout the task (**Fig. 1A**), participants fixated at a central square fixation point (0.2°), and all stimuli appeared on a neutral gray background. On each trial, we presented participants with two small colored dot stimuli (red or blue; 0.15°) as possible WM targets. After 500 ms, the targets disappeared and the fixation point changed color to red or blue to act as a post-cue which reliably informed participants which of the two positions to maintain in visual spatial WM over an extended delay period. After a 16,000 ms delay period, a response bar appeared on the screen (randomly oriented horizontally or vertically) and participants adjusted the position of the response bar using a button box held in their right hand to match the corresponding coordinate of the remembered position (e.g., if the bar was horizontal, the buttons moved the bar up/down, and the participants reported the y coordinate of the remembered position). Participants had 3,000 ms to adjust the bar to make their response, and we took the final position of the bar as the reported location on that trial. WM targets could appear within 2.9-4.1° eccentricity and at random polar angles and were presented at either 72° or 144° polar angle apart, with ∼10° polar angle jitter (counterbalanced).

**Figure 1:**
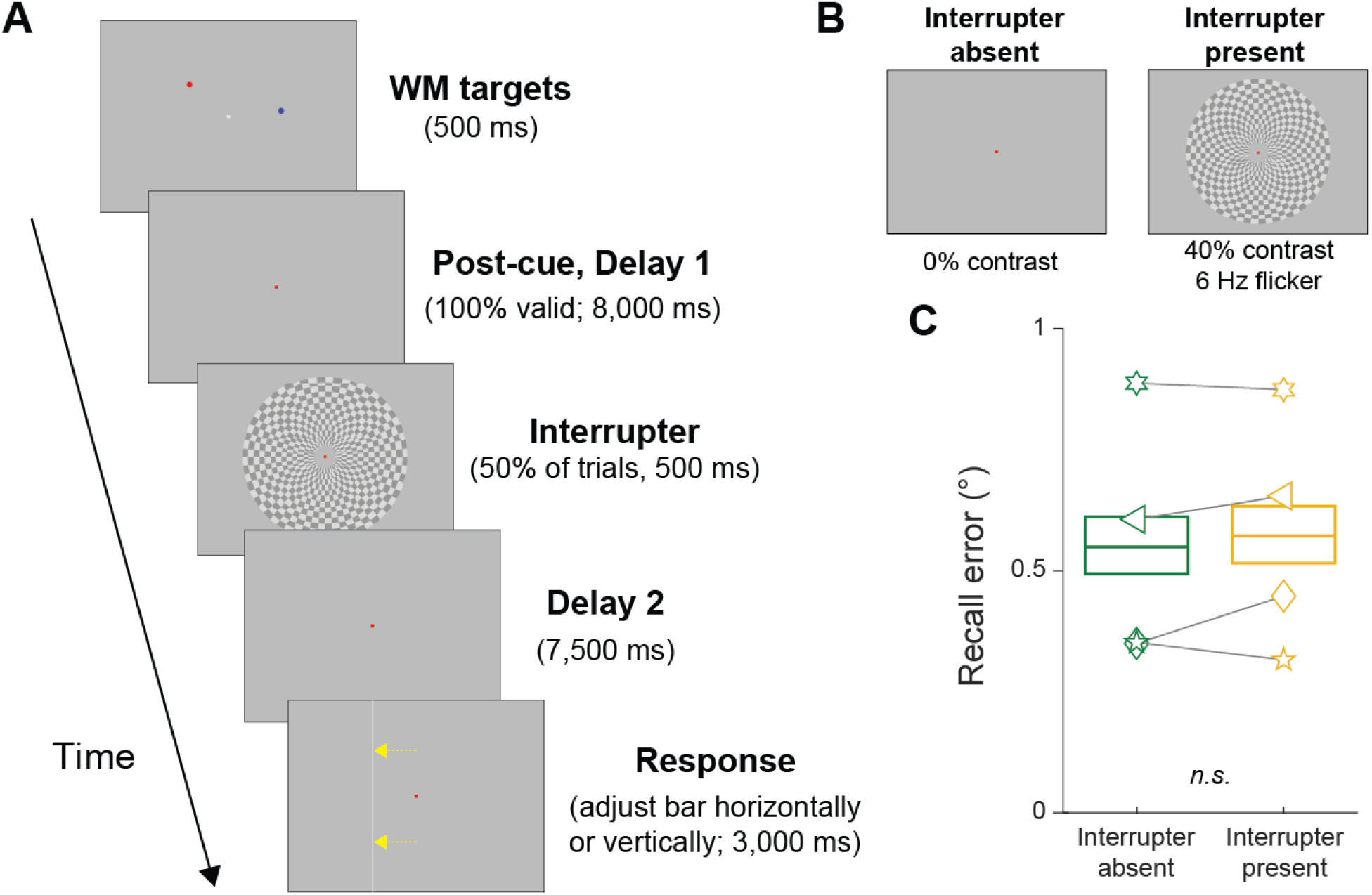
Spatial working memory task with intermittent visual disruption. (**A**) On each fMRI trial, participants (n = 4; 2x 2-hr scanning sessions each) viewed two dots (red and blue) presented ∼3.5° from fixation at pseudorandom locations on the screen. Immediately after the dots disappeared, we post-cued participants which of the two dots to remember for a 16 s delay period. At the end of the delay period, participants reported either the horizontal or vertical position (randomly selected on each trial) of the remembered target dot. (**B**) We tested two conditions: on interrupter-present trials (50%), a brief, task-irrelevant disruptive visual stimulus appeared midway through the delay. On interrupter-absent trials (50%), no intervening stimulus occurred during the delay period. (**C**) Behavioral recall performance was unperturbed by the disruptive visual stimulus (p = 0.62; randomization test; see methods). Bounds of box show 95% confidence interval around resampled mean across trials; horizontal bar in box shows mean. Individual data points are means of individual participants.

On 50% of trials a 500 ms interrupting stimulus appeared midway through the delay period (onset 8,000 ms after delay period began; **Fig. 1B**). The interrupting stimulus was a 6 Hz contrast-reversing radial checkerboard (radius: 6°; broken into 15 equal-sized radial and 60 tangential segments) presented at 40% contrast. Other than this brief interrupting stimulus trials were identical between conditions. Intertrial intervals (ITIs) were uniformly sampled between 2 and 6 s separated each trial.

We counterbalanced trials across groups of 4 6-min scanning runs, with each individual run containing 15 trials. Across a set of 4 runs, we included 3 trials each of all possible combinations of mean stimulus location (5 bins), WM target separation distance (72° or 144°; 2 possibilities), and occurrence of interrupting stimulus (present or absent; 2 possibilities). Each run lasted 366 s, including a 12 s blank period at the start of each run and a 10 s blank period at the end of each run.

### Task design: spatial mapping task

To estimate a spatial encoding model for each voxel, we acquired fMRI data while participants fixated a fixation point (0.2°) and covertly attended flickering checkerboard stimuli presented at a triangular grid of locations on the screen (**Fig. 3A**; as in previous reports; Sprague et al., 2018, 2016). Participants attended these stimuli to identify rare target events (changes in checkerboard contrast on 10 of 47 [21.2%] of trials, evenly split between increments & decrements). During each run, we chose the position of each trial’s checkerboard (0.9° radius, 70% contrast, 6 Hz full-field flicker) from a triangular grid of 37 possible positions and added a random uniform circular jitter (0.5° radius). We rotated the “base” position of the triangular grid on each scanner run to increase the spatial sampling density. Accordingly, every mapping trial was unique. The base triangular grid of stimulus positions separated stimuli by 1.5°, and extended 4.5° from fixation (3 steps). This, combined with random jitter and the radius of the mapping stimulus, resulted in a visual “field of view” (region of the visual field stimulated) of 5.9° from fixation for our spatial encoding model. On trials in which the checkerboard stimulus overlapped the fixation point, we drew a small aperture around fixation (0.8° diameter).

Each trial consisted of a 3,000 ms stimulus presentation period followed by a 2,000-6,000 ms ITI (uniformly sampled). On trials with targets, the checkerboard was dimmed or brightened for 500 ms, beginning at least 500 ms after stimulus onset and ending at least 500 ms before stimulus offset. We instructed participants to only respond if they detected a change in checkerboard contrast and to minimize false alarms. We discarded all target-present trials when estimating spatial encoding models. To ensure participants performed below ceiling we adjusted the task difficulty between each mapping run by changing the percentage contrast change for target trials. Each run consisted of 47 trials (10 of which included targets), a 12 s blank period at the beginning of the run, and a 10 s blank period at the end of the run, totaling 352 s.

### Task design: spatial localizer

To focus our neuroimaging analyses to voxels responsive during spatial WM maintenance over the area subtended by our display setup, we scanned each participant on several runs of a visual spatial WM localizer task. For some participants, this localizer was part of data acquired for a previously-reported study. On each trial we presented a flickering radial checkerboard annulus in one visual hemifield extending from 0.8° to 6.0° from fixation (1.25 cycles/° from fixation, 12° per polar angle cycle, 6 Hz contrast-reversing) for 10 s. During the stimulus interval, we presented 2 spatial WM trials in which participants remembered the precise position of 1 red dot over a 3 s delay interval. At the end of each delay interval, participants responded whether a green probe stimulus was to the left or to the right, or above or below, the remembered target position as indicated by a horizontal or vertical bar at fixation, respectively. WM targets could only appear within the stimulated hemifield. We maintained performance at ∼75% by adjusting the task difficulty (target/probe separation distance) across trials. Stimulus epochs were separated by 3 – 5 s ITIs (uniform distribution). Each run contained 4 null trials that were the same duration as normal trials but did not contain checkerboard stimuli.

### fMRI acquisition

We scanned all participants on a 3 T research-dedicated GE MR750 fMRI scanner at UC San Diego’s Keck Center for Functional Magnetic Resonance Imaging with a 32 channel send/receive head coil (Nova Medical, Wilmington, MA). We acquired functional data using a gradient echo planar imaging (EPI) pulse sequence (19.2 × 19.2 cm field of view, 64 × 64 matrix size, 35 3-mm-thick slices with 0-mm gap, axial orientation, TR = 2,000 ms, TE = 30 ms, flip angle = 90°, voxel size 3 mm isotropic).

To anatomically coregister images across sessions, and within each session, we also acquired a high resolution anatomical scan during each scanning session (FSPGR T1-weighted sequence, TR/TE = 11/3.3 ms, TI = 1,100 ms, 172 slices, flip angle = 18°, 1 mm^3^ resolution). For all sessions but one, anatomical scans were acquired with ASSET acceleration. For the remaining session, we used an 8 channel send/receive head coil and no ASSET acceleration to acquire anatomical images with minimal signal inhomogeneity near the coil surface, which enabled improved segmentation of the gray-white matter boundary. We transformed these anatomical images to Talairach space and then reconstructed the gray/white matter surface boundary in BrainVoyager 2.6.1 (BrainInnovations, The Netherlands) which we used for identifying ROIs.

### fMRI preprocessing

We coregistered functional images to a common anatomical scan across sessions (used to identify gray/white matter surface boundary as described above) by first aligning all functional images within a session to that session’s anatomical scan, then aligning that session’s scan to the common anatomical scan. We performed all preprocessing using FSL (Oxford, UK) and BrainVoyager 2.6.1 (BrainInnovations). Preprocessing included unwarping the EPI images using routines provided by FSL, then slice-time correction, three-dimensional motion correction (six-parameter affine transform), temporal high-pass filtering (to remove first-, second- and third-order drift), transformation to Talairach space (resampling to 2×2×2 mm resolution) in BrainVoyager, and finally normalization of signal amplitudes by converting to Z-scores separately for each run using custom MATLAB scripts. We did not perform any spatial smoothing beyond the smoothing introduced by resampling during the co-registration of the functional images, motion correction and transformation to Talairach space. All subsequent analyses were computed using custom code written in MATLAB (release 2015a).

### Region of Interest definition

We identified 10 *a priori* ROIs using independent scanning runs from those used for all analyses reported in the text. For retinotopic ROIs (V1-V3, hV4, V3A, IPS0-IPS3), we utilized a combination of retinotopic mapping techniques. Each participant completed several scans of meridian mapping in which we alternately presented flickering checkerboard “bowties” along the horizontal and vertical meridians. Additionally, each participant completed several runs of an attention-demanding polar angle mapping task in which they detected brief contrast changes of a slowly-rotating checkerboard wedge. We used a combination of maps of visual field meridians and polar angle preference for each voxel to identify retinotopic ROIs (Engel et al., 1994; Swisher et al., 2007). We combined left- and right-hemispheres for all ROIs, as well as dorsal and ventral aspects of V2 and V3 for all analyses by concatenating voxels.

We defined superior precentral sulcus (sPCS), the putative human homolog of macaque frontal eye field (FEF; (Mackey et al., 2017; Srimal and Curtis, 2008) by plotting voxels active during either the left or right conditions of the localizer task described above (FDR corrected, *q* = 0.05) on the reconstructed gray/white matter boundary of each participant’s brain and manually identifying clusters appearing near the superior portion of the precentral sulcus, following previous reports (Srimal and Curtis, 2008). Though retinotopic maps have previously been reported in this region (Hagler and Sereno, 2006; Mackey et al., 2017), we did not observe topographic visual field representations using our rotating wedge mapping stimulus. We anticipate this is due to the substantially limited visual field of view we could achieve inside the scanner (maximum eccentricity: ∼7°), and because we did not sample across different eccentricities.

We selected voxels for further analysis using the same procedure used to identify sPCS defined above.

### fMRI analyses: univariate

To test whether the interrupter stimulus resulted in an evoked fMRI BOLD response in visual, parietal, and frontal cortex, we computed event-related averages of each trial for each voxel, then averaged resulting timecourses across all voxels within each ROI for each trial, then across all trials within each condition. For this and all subsequent analyses, we only included voxels activated by the spatial localizer task (see above). To compute 95% confidence intervals (CIs; **Fig. 2**), we resampled all trials from all participants with replacement 1,000 times, averaged the resampled trials within each iteration, and used the 2.5^th^ and 97.5^th^ percentile of the resulting distribution as the 95% CIs.

**Figure 2:**
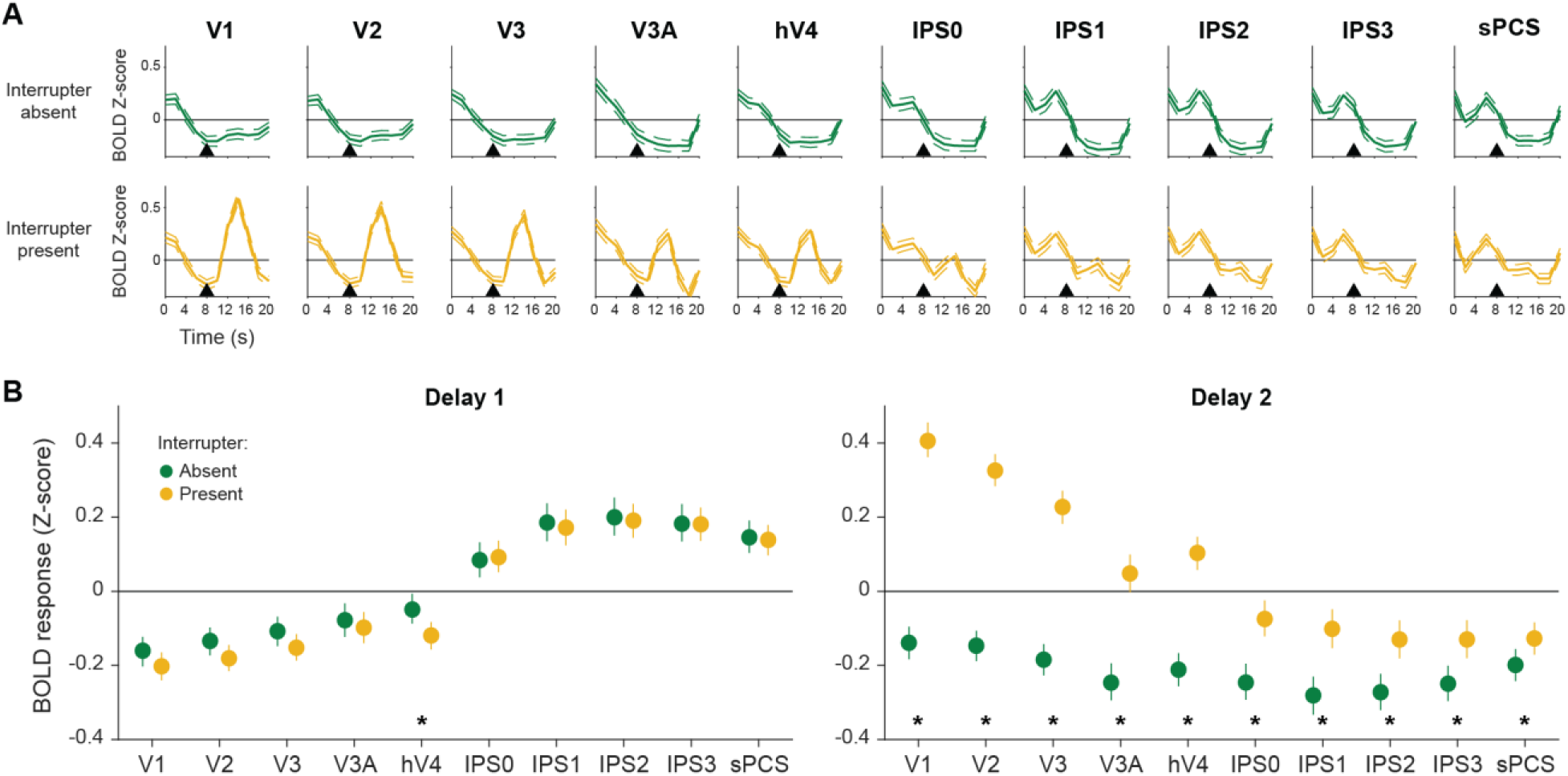
Visual regions show transient visually-evoked response to intervening visual stimulus. (**A**) Average hemodynamic response function (HRF) for each retinotopic ROI, averaged across active voxels and participants. Error band is 95% confidence interval computed via resampling across trials. Black triangle indicates onset of intervening visual stimulus (on interrupter-present trials). (**B**) Comparison of mean delay-period activation for Delay 1 (defined as TRs beginning at 6 and 8 s after delay onset) and Delay 2 (TRs beginning at 14 s and 16 s). Black asterisk indicates significant difference between conditions (q = 0.05, corrected via FDR across all comparisons); gray asterisk indicates trend (alpha = 0.05, uncorrected).

### fMRI analyses: inverted encoding model

To reconstruct spatial representations of the content of visual WM carried by activation patterns measured over entire regions of interest, we implemented an inverted encoding model (IEM) for spatial position (Sprague and Serences, 2013). This analysis involves first estimating an encoding model (sensitivity profile over the relevant feature dimension(s) as parameterized by a small number of modeled information channels) for each voxel in a region using a “training set” of data reserved for this purpose (four spatial mapping runs). Then, the encoding models across all voxels within a region are inverted to estimate a linear mapping that can be used to transform novel activation patterns from a “test set” (spatial WM task runs) into activation in the modeled set of information channels.

We built an encoding model for spatial position based on a linear combination of spatial filters (Sprague et al., 2016, 2014; Sprague and Serences, 2013). Each voxel’s response was modeled as a weighted sum of 37 identically-shaped spatial filters arrayed in a triangular grid (**Fig. 3B**). Centers were spaced by 2.27° and each filter was a Gaussian-like function with full-width half-maximum of 2.49°:

**Figure 3:**
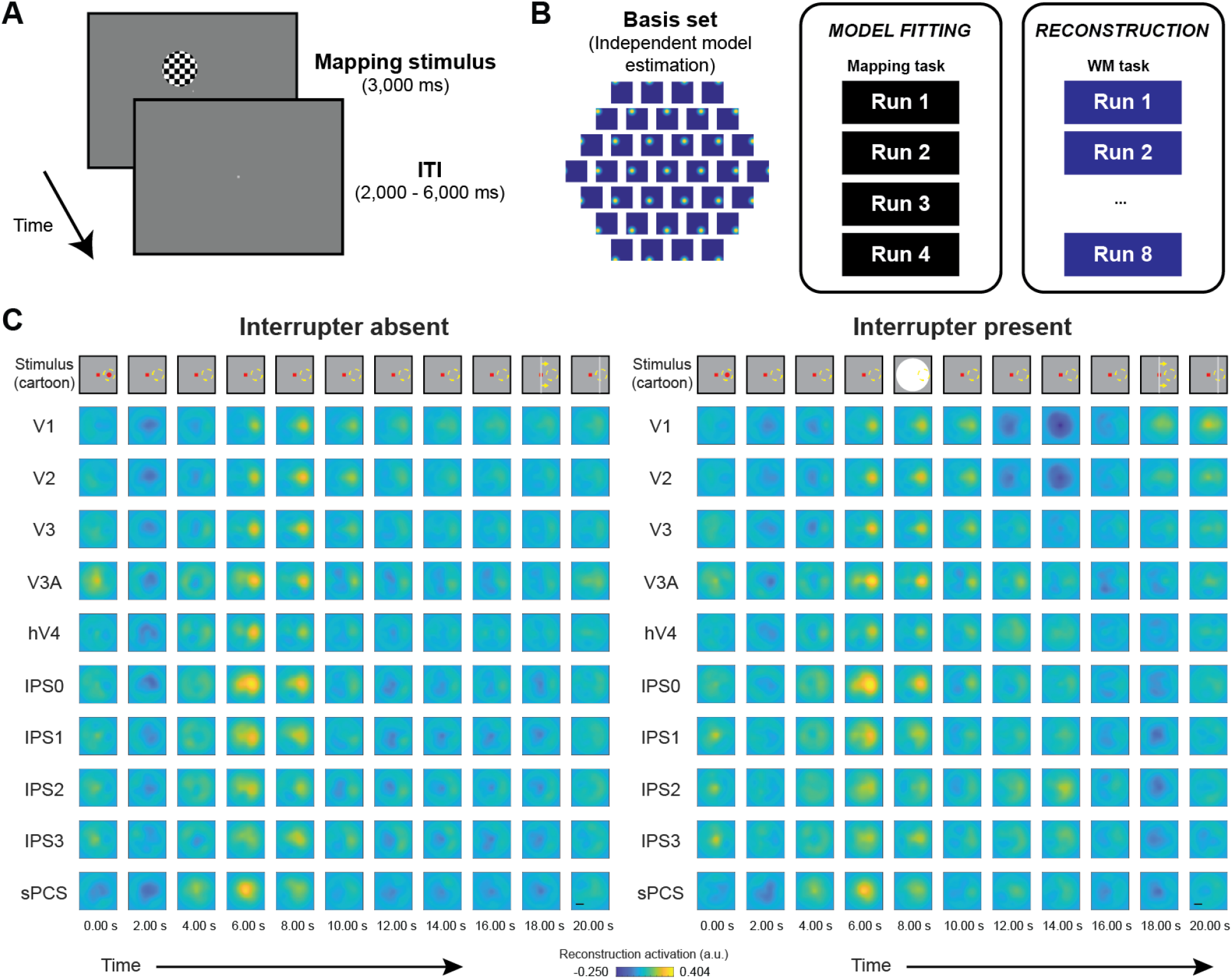
Model-based spatial reconstructions of WM representations exhibit global activation following interruption. (**A**) During each session, participants completed several runs of a ‘mapping’ task used to fit an inverted encoding model (IEM) of visual spatial location. On each trial, participants attended to a flickering checkerboard presented at a pseudorandom location on the screen and reported rare contrast increments/decrements (22% of trials). The stimulus appeared at a jittered location drawn from a triangular grid. (**B**) The IEM comprised a basis set of 37 spatially-selective channels (e.g., modeled neural populations with spatially-selective receptive fields) defined as a triangular grid. To fit the model (see Methods), we estimated the contribution of each channel to each voxel as the best-fit linear combination of predicted responses from each channel on each trial of the mapping task. Then, we used the best-fit model to reconstruct each trial and timepoint of the WM task (**Fig. 1A**). See Methods for details of model-fitting and reconstruction procedures. This “fixed encoding model” approach (Sprague et al., 2018a) ensures all trials and timepoints are reconstructed on equal footing and can be directly compared with one another. (**C**) Spatial reconstructions of the visual field, aligned to the remembered spatial location on each trial and averaged across trials and participants. All images plotted on the same color scale to allow for direct comparison between panels. WM location aligned to the horizontal positive axis. On interrupter-absent trials, spatial WM representations are visible throughout the delay period, though are substantially weakened at later points in the delay period (∼14-16 s). On interrupter-present trials in which an intervening visual stimulus appears midway through the delay period, reconstructions exhibit a global change associated with the transient visually-evoked response (**Fig. 2**). While this effect is variable across ROIs, the WM representation is still present within reconstructions. See **Fig. 4** for quantitative comparison of these reconstructions. Scale bar 3°. Top line is cartoon depiction of (aligned) stimulus/remembered locations (red dot and yellow dashed circle, respectively), not shown to scale. Interrupter stimulus was a flickering radial checkerboard, as shown in **Fig. 1A-B**.

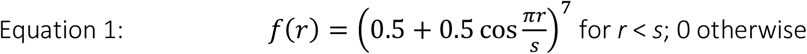

Where *r* is the distance from the filter center and *s* is a “size constant” reflecting the distance from the center of each spatial filter at which the filter returns to 0. Values greater than this are set to 0, resulting in a single smooth round filter at each position along the triangular grid (*s* = 6.27°; see **Fig. 3B** for illustration of filter layout and shape).

This triangular grid of filters forms the set of information channels for our analysis. Each mapping task stimulus is converted from a contrast mask (1’s for each pixel subtended by the stimulus, 0’s elsewhere) to a set of filter activation levels by taking the dot product of the vectorized stimulus mask and the sensitivity profile of each filter. This results in each mapping stimulus being described by 37 filter activation levels rather than 1,024 × 768 = 786,432 pixel values. Once all filter activation levels are estimated, we normalize so that the maximum filter activation is 1.

We model the response in each voxel as a weighted sum of filter responses (which can loosely be considered as hypothetical discrete neural populations, each with spatial RFs centered at the corresponding filter position).

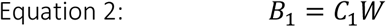

Where *B*_*1*_ (*n* trials × *m* voxels) is the observed BOLD activation level of each voxel during the spatial mapping task (averaged over two TRs, 6.00-8.00 s after mapping stimulus onset), *C*_*1*_ (*n* trials × *k* channels) is the modeled response of each spatial filter, or information channel, on each non-target trial of the mapping task (normalized from 0 to 1), and *W* is a weight matrix (*k* channels × *m* voxels) quantifying the contribution of each information channel to each voxel. Because we have more stimulus positions than modeled information channels, we can solve for *W* using ordinary least-squares linear regression:

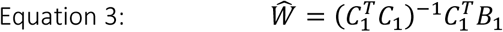

This step is univariate and can be computed for each voxel in a region independently. Next, we used all estimated voxel encoding models within a ROI (***Ŵ***) and a novel pattern of activation from the WM task (each TR from each trial, in turn) to compute an estimate of the activation of each channel (***Ĉ***_**2**_, *n* trials × *k* channels) which gave rise to that observed activation pattern across all voxels within that ROI (*B*_*2*_, *n* trials × *m* voxels):

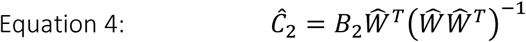

Once channel activation patterns are computed (Equation 4), we compute spatial reconstructions by weighting each filter’s spatial profile by the corresponding channel’s reconstructed activation level and summing all weighted filters together. This step aids in visualization, quantification, and coregistration of trials across stimulus positions, but does not confer additional information. Because stimulus positions were unique on each trial of the spatial WM task, direct comparison of image reconstructions on each trial is not possible without coregistration of reconstructions so that stimuli appeared at common positions across trials. To accomplish this, we adjusted the center position of the spatial filters on each trial such that we could rotate the resulting reconstruction. For **Figure 3**, we rotated each trial such that the remembered target was centered at *x* = 3.5° and *y* = 0°.

In our primary analyses (**Figs. 3-4**), we used 4 mapping task runs to estimate the encoding model for each voxel, then inverted that encoding model to reconstruct visual field images during all main spatial attention task runs (**Fig. 3B**). In a subsequent analysis we estimated encoding models using all runs but one for each timepoint and condition of the experiment (**Fig. 5**).

**Figure 4:**
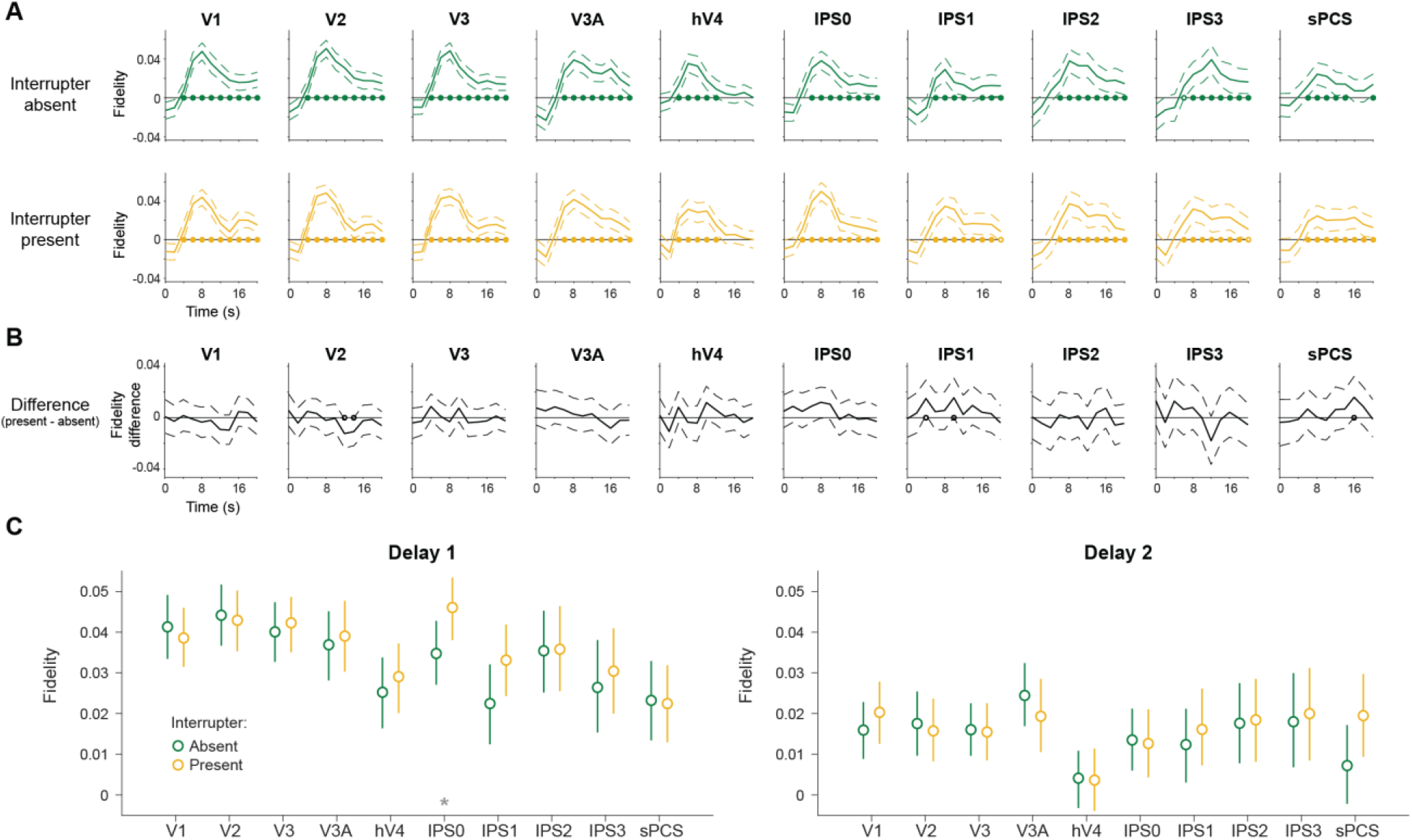
WM representation fidelity does not change following visual interruption. (**A**) Timecourse of WM representation fidelity (see Methods) computed at each timepoint during the delay. There is no apparent difference between WM representation fidelity on interrupter-absent and -present trials. Solid circles indicate timepoints with significant fidelity based on a resampling test (q = 0.05, corrected via FDR across all timepoints, conditions, and ROIs); open circles indicate timepoints with trends (alpha = 0.05, no correction for multiple comparisons). Error bands 95% CI based on resampling across trials. **(B)** Difference timecourse of WM representation fidelity (interrupter present – interrupter absent). Solid/open circles indicate significant differences/trends between conditions, as defined in (A). Error bands 95% CI. **(C)** Comparison of fidelity between conditions during Delay 1 and Delay 2 for each ROI. Each point shows the average fidelity computed across the Delay 1 and Delay 2 epochs, respectively, and error bars are 95% CIs based on resampling across trials. Black asterisks indicate significant difference between conditions corrected via FDR across all ROIs and both delays, q = 0.05; gray asterisks indicate trends (alpha = 0.05, no correction for multiple comparisons).

**Figure 5:**
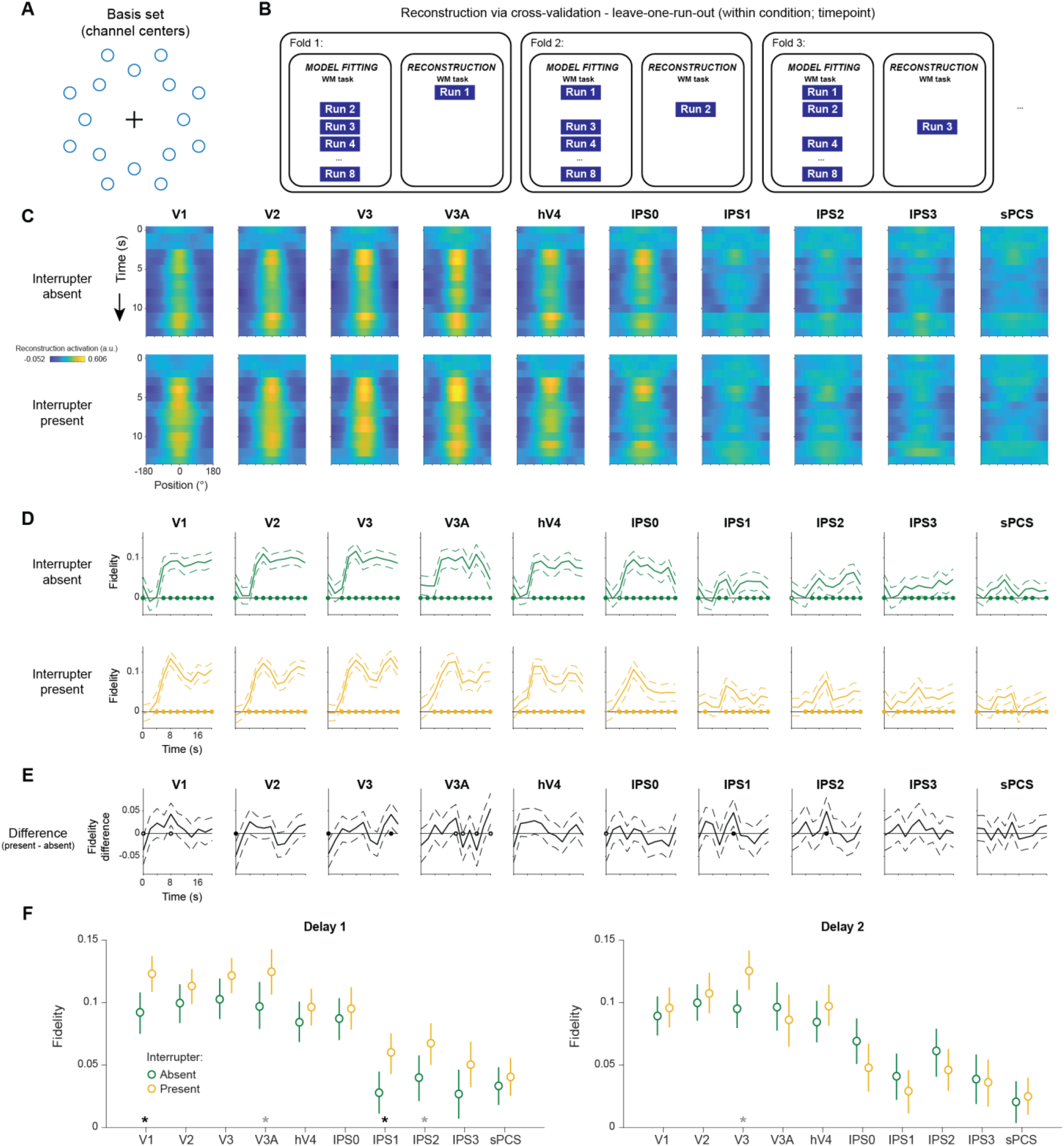
Matched model estimation does not reveal evidence for a morphed code following visual stimulation. **(A)** In this analysis, we implemented leave-one-run-out (LORO) cross-validation and estimated a modified inverted encoding model using each timepoint’s activation associated with each condition. Because WM locations only spanned an anulus between 2.9° and 4.1° eccentricity, we computed a reduced IEM spanning that range, with channels centered along two iso-eccentric rings (see Methods). **(B)** We implemented our LORO procedure for each condition and trial timepoint separately. We estimated an IEM using the reduced basis shown in (A) using all runs but one and inverted this model to reconstruct spatial representations on matched trials of the held-out run. While this does not allow for direct comparison of data from each condition on equal footing, this procedure is in principle more sensitive to any differences in the format of WM representations across time and between conditions. **(C)** Comparison of spatial reconstructions (channel activation as a function of polar angle relative to remembered position) through time for each condition and ROI. All panels are plotted on the same color scale to allow for direct comparison between conditions and ROIs. **(D)** Quantification of WM representational fidelity computed using reconstructions in (C). Filled/open circles indicate statistical differences as described in **Fig. 4A**. Error bands show 95% CIs computed via resampling. **(E)** Difference timecourse computed between stimulus-present and -absent conditions (as in **Fig. 4B**). Filled/open circles indicate statistical differences as described in **Fig. 4B**. Error bands show 95% CIs of difference timecourse computed via resampling. **(F)** Comparison of fidelity between conditions during Delay 1 and Delay 2 for each ROI, as in **Fig. 4C**. Each point shows the average fidelity computed across the Delay 1 and Delay 2 epochs, respectively, and error bars are 95% CIs based on resampling across trials. Black asterisks indicate significant difference between conditions corrected via FDR across all ROIs and both delays; gray asterisks indicate trends (alpha = 0.05, no correction for multiple comparisons).

In analyses in which we performed leave-one-run-out cross-validation (**Fig. 5**), we used a different spatial arrangement of channels as our basis set. This was required because remembered targets never appeared more foveal than 2.9° eccentricity, and so it would be impossible to reliably estimate the contribution of channels positioned within that area. Instead, we used a staggered ring of spatial channels (see **Fig. 5A**) centered at 2.9° and 4.1° eccentricity, and each separated by 45° deg polar angle and interdigitated (inner ring 0°, 45°, etc; outer ring 22.5°, 67.5°, etc). Additionally, rather than estimate a model using entirely separate ‘mapping’ data, we estimated the model using all runs except for one of each session’s WM dataset and used this model to reconstruct channel responses from the held-out run (**Fig. 5B**). Moreover, we employed a matched training/testing procedure whereby each reconstruction was computed with a model estimated using the corresponding timepoint and task condition (interrupter present/absent). This, in principle, should ensure the highest chance of observing a WM representation should it be present, and should guard against the possibility of observing lower-fidelity reconstructions due to a mismatch between the coding format of the WM representation and that estimated using separate training data (as in **Fig. 3**).

### Quantifying WM representations: fidelity

To quantify spatial WM representations in reconstructions, we defined a “representational fidelity” metric that quantifies the extent to which a target representation reliably appeared within a reconstruction (Rademaker et al., 2019; Sprague et al., 2016). To accomplish this, we first reduced the reconstruction from a 2-d image to a 1-d line plot by averaging over each of 220 evenly-spaced polar angle arms subtending 2.9-4.1° eccentricity. The resulting 1-dimensional reconstruction reflects the average profile along an annulus around fixation. A target representation in these reconstructions would be a “bump” near 0° after the reconstructions have been rotated to a common center (where 0° corresponds to the actual target polar angle). To reduce these 1-d reconstructions to a single number which could be used to quantify the presence of target representations (*F*), we computed a vector mean of the 1-d reconstruction (*r*(*θ*), where *θ* is the polar angle of each point and *r*(*θ*) is the reconstruction activation) when plotted as a polar plot, as projected along the *x* axis (because the reconstructions were rotated such that the target was presented at 0°):

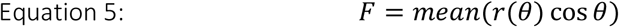

If *F* is reliably greater than zero, over a resampling procedure (see Statistical Procedures), this quantitatively demonstrates that the net activation over the entire reconstruction carries information above chance about the target position. This measure is independent of baseline activation level in the reconstruction, as the mean of *r*(*θ*) is removed by averaging over the full circle. We computed timecourses of representational fidelity (**Fig. 4A-B**), as well as representational fidelity for each delay period (**Fig. 4C**). To determine whether the interrupting stimulus on interrupter-present trials impacts WM representations, we compared *F* between Delay 2 and Delay 1 for each ROI.

### Statistical procedures

All statistical statements reported in the text are based on resampling procedures in which a variable of interest is computed over 1,000 iterations. In each iteration, all single-trial variables from a given condition are resampled with replacement and averaged, resulting in 1,000 resampled averages for a given condition. We then subjected these distributions of resampled averages to pairwise comparisons by computing the distribution of differences between one resampled distribution (e.g., interrupter absent) and another resampled distribution (e.g., interrupter present), yielding a new distribution of 1,000 difference values. We tested whether these difference distributions significantly differed from 0 in either direction by performing two one-tailed tests (*p* = proportion of values greater than or less than 0; null hypothesis that difference between conditions = 0) and doubling the smaller *p* value. For tests in which we compared whether representations were present in 1-d reconstructions using the representational fidelity measure, we performed one-tailed tests (null hypothesis that *F* ≤ 0).

Because we performed 1,000 iterations of these analyses, we cannot identify *p* values less than 0.001, so all comparisons in which resampled difference distributions were all greater than or less than 0 are reported as *p* < 0.001. Because we performed many pairwise comparisons, we corrected all repeated tests within an analysis using the false discovery rate (Benjamini and Yekutieli, 2001) and a threshold of *q* = 0.05. All error bars/intervals reflect 95% confidence intervals as estimated using this resampling procedure.

### Data and code availability

[Upon publication of this report] all data and code [will be] made freely available on the authors’ GitHub and Open Science Framework pages. Available data includes all files necessary to produce figures in this report: behavioral data from fMRI scanning sessions, and extracted data from each ROI on each timepoint of each trial.

## Results

### Behavioral recall performance

In this experiment, participants (n = 4) were asked to report the horizontal or vertical location of one of two dots after a 16 second delay (**Fig. 1A**). On 50% of trials, a brief task irrelevant stimulus was presented during the delay (interrupter-present), while on the other 50% of trials, the blank screen remained unchanged throughout the 16 s delay period (interrupter-absent; **Fig. 1B**). The intervening stimulus did not significantly impact the accuracy of the reports (interrupter-present vs interrupter-absent, p = 0.62, resampling test; see Materials & Methods; **Fig. 1C**).

### fMRI results: univariate

We computed univariate bold responses by averaging response timecourses across all voxels within each ROI. First, a direct comparison of event-related averaged timeseries (**Fig. 2A**) reveals an overall decrease in average activation during the WM delay period in early visual regions, and a transient increase in average activation in retinotopic parietal regions, likely associated with attending to and encoding the remembered location. Following the irrelevant visual stimulus, we observed a transient evoked response that was especially strong in early visual ROIs and was more measured in retinotopic parietal regions. Qualitatively, these results are consistent with previous observations that single remembered spatial locations do not always result in sustained activation across all voxels in visual, parietal, or frontal cortex (Sprague et al., 2016, 2014; but see Curtis and Sprague, 2021), and demonstrate the effectiveness of our interrupter stimulus at driving visually-evoked cortical activity.

To quantify whether average BOLD activation differed between interrupter-absent and interrupter-present conditions, we averaged activation over each delay period (Delay 1: 6 to 10 s; Delay 2: 14 to 18 s), then, within each delay period, resampled all trials across all participants with replacement, computed a difference score, and compared the distribution of difference scores against 0 (**Fig. 2B**). Activation measured during Delay 1 was indistinguishable between conditions in all ROIs except for hV4 (p = 0.01; resampling test, corrected for multiple comparisons via false discovery rate [FDR]). This difference is likely to be spurious, as no aspect of the task or stimulus differed between condition at this stage of the trial. During Delay 2, there is significantly greater BOLD activation in visual areas following the presentation of the irrelevant stimulus when compared to interrupter-absent trials in all ROIs (maximum p = 0.022, sPCS; **Fig. 2B**). This confirms that the task-irrelevant stimulus resulted in greater BOLD activation throughout the visual system.

However, from these univariate analyses alone, it remains unclear whether the fidelity of working memory representations carried by activation patterns were significantly affected by the intervening stimulus. Previous studies have found evidence that, under different circumstances, working memory representation fidelity can be degraded (Bettencourt and Xu, 2016), biased (Hallenbeck et al., 2021; Lorenc et al., 2018), augmented (Wolff et al., 2017, 2015), left intact (Rademaker et al., 2019) or transformed (Parthasarathy et al., 2017) by intervening interrupting stimuli. Next, to quantitatively compare the fidelity of information carried by activation patterns in retinotopic ROIs, we employed a multivariate model-based analysis approach which allowed us to determine whether WM representations changed significantly as a function of interrupter presentation.

### fMRI results: inverted encoding model

We implemented an inverted encoding model (IEM) to decode the contents of working memory. Briefly, this model allows us to reconstruct the spatial representation supported by the activation pattern across a region at each time point of the experiment. First, we fit an encoding model based on a discrete number of neural channels, which can be thought of as modeled spatially selective neural populations, each channel being selective for a different location. The response of each voxel is modeled as a weighted sum of these channel responses. Using a set of data independent from the main task (**Fig. 3A-B**), we can measure how each voxel responds to various stimuli presented at different locations on the screen. Using this independent fMRI data along with the predicted channel responses, we can use linear regression to estimate a set of weights for each channel within each voxel, which acts as our encoding model. Next, we invert this model to reconstruct channel activation based on observed BOLD activation patterns in each ROI during the WM task. Finally, we compute a weighted sum of the modeled channel profiles weighted by the reconstructed channel activation, and rotate and align the resulting reconstructed spatial representations of the modeled visual field such that the remembered location is at a fixed and known position. This implementation whereby we estimate a model using an independent dataset is an example of a ‘fixed encoding model’ approach, and ensures that comparisons between timepoints and conditions are on equal footing because the same model is used to compute reconstructions that are directly compared to one another (Sprague et al., 2019, 2018a).

Stronger activation within reconstructions aligned with the remembered position, indicated by brighter colors, are due to activation patterns during the delay period supporting a spatial WM representation of the remembered position (**Fig. 3C**). Model-based reconstructions computed from each trial are rotated to align all the remembered locations to a single location, because the particular remembered location on each trial is not of interest. Lastly, all aligned trials over all participants are averaged together at each timepoint (**Fig. 3C**). First, focusing on the interrupter-absent reconstructions (**Fig. 3C**; left), there is a peak in the reconstruction strength in all ROIs at approximately 6-8 s after target encoding (note that these timecourses are not corrected for the delay in the hemodynamic response). These representations lose strength through the delay period, but nevertheless exist until the participant makes their behavioral response (16 s). Next, turning to the interrupter-present trials, if a brief interrupter can cause a decrease in the strength of the WM representation, we would expect WM representations to disappear from reconstructions following the interrupter presentation. Alternatively, if the intervening visual stimulus leads to an increase in local processing that augments WM representations, we would expect to see a transient strengthening of the WM representation in reconstructions above a noisy baseline. However, **Fig. 3C** shows that WM representations within reconstructions appear to persist throughout the delay period, and are not obviously disturbed nor augmented by the irrelevant visual stimulation. While this qualitative result is consistent with the possibility that WM representations are robust to visual interruption, it is difficult to discern how much the strength of these representations change, if at all, based on visual inspection alone.

We quantified the quality of WM representations in these reconstructions using a ‘fidelity’ metric, which measures how much information there is about the remembered position (see Materials & Methods). This is accomplished by representing each location of the reconstruction as its polar angle from the center, and measuring the amount of reconstruction activation for each polar angle, represented as a vector on a circle. Then, every vector is averaged together, and the resulting mean vector is projected onto a unit vector pointing in the direction of the remembered position (Rademaker et al., 2019; Sprague et al., 2016). If there is a reliable representation of the remembered position within a reconstruction, this fidelity metric should be greater than 0. Looking at both the interrupter-absent and the interrupter-present trials, consistent with the working memory reconstructions, there is a reliable representation throughout the delay period in each ROI (**Fig. 4A**), with the exception of hV4 (one-tailed resampling test against 0; FDR-corrected across all ROIs, timepoints, and conditions; q = 0.05). Critically, we found no difference between the WM representation fidelity on interrupter-absent and interrupter-present trials at any timepoint (**Fig. 4B**; two-tailed resampling test against 0, FDR-corrected across all ROIs and timepoints; q = 0.05).

Next, we averaged reconstructions over Delay 1 and over Delay 2 and, within each delay period, directly compared fidelity between interrupter-absent and interrupter-present trials (**Fig. 4C**). During Delay 1, we found a trending effect wherein the WM representation fidelity was greater during interrupter-present compared to interrupter-absent trials in IPS0 (p = 0.044; two-tailed resampling test against 0, not corrected for multiple comparisons). No other regions showed a significant difference nor trend during either delay period (minimum p = 0.072, sPCS, Delay 2). Similar to the difference seen in hV4 in the univariate analyses, we believe this result is likely spurious, because the experimental procedure up until the end of Delay 1 is identical for both conditions. Combined with the univariate analyses, we have evidence to conclude that working memory representations in primary visual cortex and IPS remain stable following irrelevant visual information.

The multivariate analyses employed thus far involved estimating an encoding model using an entirely independent dataset which required sustained spatial attention to visually-presented checkerboard discs (Sprague et al., 2018a). Our results revealed no significant change in WM representation fidelity following visual interruption (**Fig. 4**). However, it may be the case that the WM representation is in fact disrupted (or augmented), just not in a format that’s accessible via our independently-trained model. For example, the brief visual stimulus could result in a morphed neural code (Parthasarathy et al., 2017), and so differences in WM representations would be observed if a model was estimated using the same experimental data (at the same timepoints, and within the same conditions) as used for reconstruction. Indeed, a recent report illustrates the importance of comparing results between model estimation procedures by showing that results can qualitatively change when different datasets are used to train a decoding algorithm (Iamshchinina et al., 2021). To test this possibility, we implemented a leave-one-run-out (LORO) cross-validation approach. In the LORO procedure, we estimate the encoding model for each timepoint and condition using data from all runs of the WM task except for one. Then, we reconstruct channel activation profiles using data from this held-out run, and repeat so that all trials are reconstructed (see Methods; **Fig. 5B**). Note that because WM stimuli only appeared within an anulus, we could only estimate a spatial model for a fixed eccentricity range (**Fig. 5A**). There are a few key differences between the two methods. First, using a fixed encoding model for reconstruction is ideal for verifying that the model is picking up on signals associated with the relevant feature dimension rather than idiosyncrasies associated with the task stimuli. In other words, if the model can successfully *generalize* from attended visual stimuli to remembered spatial positions, this means the two task domains occupy an overlapping coordinate system (Harrison and Tong, 2009). However, using an independent task for model training comes at the potential cost of decoding sensitivity. As mentioned above, if the training set and task data are well matched, the underlying neural codes will be largely generalizable from one task to the other. However, poor decoding performance may not be diagnostic of a lack of available information, since the independent training data and the task data necessarily differ in their neural codes to a greater extent than task data differs from itself across scanning runs.

The pattern of results in our study is fairly consistent regardless of analysis procedure. Like with a fixed encoding model, the WM representation persists throughout both delay periods (**Fig. 5C-D**). There are not consistent differences between the interrupter present and interrupter absent trials, but there are a few individual timepoints in which interrupter-present trials have different fidelity than interrupter-absent trials (**Fig. 5E**). However, these timepoints are isolated, and the direction of the difference is not consistent across timepoints or regions, nor is it reliably related to specific task events. Next, when we compare fidelity averaged over each delay period (**Fig. 5F**), we observed a significantly-greater fidelity on interrupter-present trials than interrupter-absent trials during Delay 1 in V1 and IPS1 (both p’s = 0.004), and trends in V3A (p = 0.026) and IPS2 (p = 0.024; both not corrected for FDR). However, because the stimulus display is identical up through this point in the trial between the two conditions, it is likely that such differences are spurious. During Delay 2, there is no difference between conditions, except a trend in the same direction in V3 (p = 0.01, not corrected for FDR; **Fig. 5F**). The direction of this trend also indicates that the interrupter present trials have stronger representations than the interrupter absent trials, consistent with a transient increase in the strength of the WM representation evoked by a ‘ping’ (e.g., Wolff et al., 2015).

Overall, between the behavioral, univariate, and multivariate results, there is little consistent evidence to suggest that working memory is susceptible to degradation due to irrelevant visual interruption, nor is there consistent evidence that a disruptive visual stimulus transiently enhances neural codes as measured with fMRI.

## Discussion

We tested whether a task-irrelevant visual stimulus presented midway through an extended delay period could impact neural activation patterns associated with visual spatial WM. If neural population codes are subject to interruption or distraction, we would expect to see a transient decrease in the quality of the WM representation during the delay period. Conversely, if activity-silent coding schemes based on changes in synaptic efficacy contribute in a substantial manner to spatial WM maintenance, we would expect to see a transient augmentation of the neural WM representation following a visual stimulus. However, we found support for neither of these alternatives. Instead, spatial WM representations were remarkably robust to visual interruption, showing neither a transient dip nor bump in coding fidelity (**Figs. 3-4**). This could not be explained by an insufficiently-stimulating interrupter stimulus, as we found positive evidence for evoked visual responses following the interrupter (**Fig. 2**), nor by a transition to an alternative coding format, as results were consistent across analyses using an independently-estimated encoding model and one estimated using cross-validation (**Fig. 5**). Finally, these results matched our observation that participants’ behavior was not impacted by the interrupting stimulus (**Fig. 1C**).

### WM representations are robust to interruption

Our observation of stable spatial WM representations in human cortex following a task-irrelevant interrupting visual stimulus is consistent with previous reports that WM representations are remarkably robust to most forms of visual ‘distraction’ (reviewed in Lorenc et al., 2021; Pasternak and Greenlee, 2005). Most experiments finding positive evidence for disruption of visual WM performance (and neural representations) involve using a distracting stimulus extremely similar to that held in WM. For example, Rademaker et al. (2019, 2015) found biased and degraded orientation recall performance when an oriented grating was used as the intervening stimulus, and this effect was particularly strong when the distracting orientation was similar to the remembered orientation. A similar effect has been observed for remembered spatial positions (Hallenbeck et al., 2021; Macoveanu et al., 2007), with behavioral performance and neural representations being impaired and/or biased when a task-relevant distracting stimulus appeared nearby the remembered position. However, none of these previous studies evaluated the neural representation of a remembered spatial position following task-irrelevant visual interruption. Our results offer evidence that spatial WM is robust to such a visual stimulus, and contribute to the ongoing discussion of whether and how neural representations are susceptible to interference from concurrent visual stimulation (Chota and Van der Stigchel, 2021; Gayet et al., 2018; Iamshchinina et al., 2021; Lorenc and Sreenivasan, 2021; Postle and Yu, 2020; Scimeca et al., 2018; Teng and Postle, 2021; Xu, 2021, 2020, 2018, 2017).

The lack of a behavioral effect of the interrupting stimulus in our experiment is important to carefully consider. Several of the previously-mentioned studies typically observed coherent changes in behavioral performance and neural representations. Lorenc et al (2018) and Hallenbeck et al (2021) both found evidence that biased neural representations in visual, but not parietal, cortex were associated with biases in behavioral performance. Rademaker et al (2019) implemented a subtle manipulation between the properties of the distracting stimuli used in their Expt 1 and Expt 2, which resulted in a significant behavioral effect of distraction in Expt 2 despite a lack of an effect in Expt 1. Interestingly, the fidelity of neural representations in visual cortex followed this pattern: when behavioral performance was adversely impacted by the interrupting stimulus in Expt 2, the neural representations were also degraded. Perhaps, then, if we had used a more efficacious interrupting stimulus – for example, a checkerboard with greater contrast – we may have seen both a decrement in behavioral performance and impaired neural WM representations. Note, though, that the specifics of the result reported in (Rademaker et al., 2019) depend on the particular analysis choices implemented (Iamshchinina et al., 2021). Here, we directly tested for this possibility by computing our results using both a ‘fixed encoding model’ (Sprague et al., 2018a) trained using an independent mapping dataset reserved for that purpose (**Figs 3-4**) and using cross-validation procedures common across machine learning techniques (**Fig. 5**). Results were consistent across these model estimation procedures.

### Activity-silent WM representations

Historically, it has been well-established that sustained delay-period activity in tuned neurons and/or voxels supports visual, and, especially, spatial, WM (Curtis and Sprague, 2021; Sreenivasan and D’Esposito, 2019). However, several recent reports have offered support for a complementary coding scheme whereby WM representations can be maintained in an activity-silent state, typically via transient changes in synaptic efficacy (Lewis-Peacock et al., 2012; Mongillo et al., 2008; Rose et al., 2016; Sprague et al., 2016; Stokes, 2015; Wolff et al., 2017, 2015). Such a coding scheme could be more metabolically efficient, as during a delay period a WM representation may be maintained by only minimal spiking, as opposed to the strong sustained spiking required by persistent activity. The most compelling evidence for this neural mechanism involves studies imposing an external stimulus – either a task-irrelevant visual flash or a TMS pulse – to ‘ping’ or ‘resurrect’ an otherwise latent neural representation (Rose et al., 2016; Wolff et al., 2020, 2017, 2015). Additional evidence for these codes derives from providing participants with an informative cue during the delay period, which results in augmentation of a weakened neural representation (Ester et al., 2018; Sprague et al., 2016; but see Schneegans and Bays, 2017). Across these studies, following the interrupting stimulus or instructive cue, the WM representation was observed to strengthen as assayed using multivariate methods similar to those used here. It stands to reason, then, that a task-irrelevant interrupting stimulus could involuntarily reinstate and/or strengthen a latent or weakened neural representation as measured with fMRI.

In our study we found no evidence for such a strengthening across multiple analysis strategies (**Fig. 3-5**). There are 3 possible explanations for our failure to find this effect. First, it may be the case that the ‘ping’ stimuli used to probe activity-silent representations primarily act to synchronize ongoing activity, which makes the activity more easily detectable at the EEG electrode on the scalp. Indeed, a recent reanalysis of prominent reports of ping-evoked neural representations suggests that there exist identifiable WM representations in patterns of alpha band activity prior to the ping (Barbosa et al., 2021; but see Wolff et al., 2021). If the trial-by-trial phase of this activity is random, but the ping serves to synchronize the ongoing oscillations, this could explain why we did not see interrupter-evoked enhancement of WM representations: the fMRI signal would not be expected to change its amplitude substantially following a shift in phase of ongoing oscillations (Hermes et al., 2017). Second, and most intuitively, it may be that our interrupter stimulus did indeed evoke a transient improvement in WM representation fidelity, but this improvement was too brief to be detected with fMRI. It is possible that future experiments implementing more rapid and/or sensitive fMRI scanning protocols, potentially in concert with simultaneous EEG measurements, could determine whether a ping-evoked strengthening of a WM representation is detectable in the BOLD signal. Third, the previous studies examining ping-evoked WM representations primarily use EEG methods, which may be limited in their signal-to-noise ratio. Perhaps the ping serves to enhance the SNR of an already-present representation so that it is detectable at the scalp. Importantly, a lack of significant decoding prior to the ping stimulus should not be taken as evidence of the absence of an active neural code, since an active neural representation may be present but undetectable using a given imaging modality (Dubois et al., 2015; Sprague et al., 2016). Altogether, we suggest that future studies using disruptive visual stimuli to test for activity-silent WM representations could be enhanced by combining multiple neuroimaging modalities to assay a full complement of temporal, spatial, and physiological resolutions (e.g., Itthipuripat et al., 2019).

### Rehearsal via spatial attention

Finally, it may be the case that the spatial WM representations decoded from fMRI activation patterns in this and previous studies are not actually due to local storage of WM information, but instead due to ongoing rehearsal processes, potentially via sustained spatial attention (Awh et al., 1998; Awh and Jonides, 2001). That is – because the location of the stimulus would covary with a spatial attention rehearsal signal, in many studies, it is impossible to dissociate these possibilities (Postle and Yu, 2020). Indeed, the spatial location of a remembered orientation or color can be decoded from EEG alpha activity patterns, even when location is irrelevant for the task at hand (Foster et al., 2017). Additionally, when spatial attention is directed to a different region of space from the remembered position, WM representations identified from fMRI activation patterns and EEG alpha patterns are transiently disrupted (Hallenbeck et al., 2021; van Moorselaar et al., 2018). Other studies using non-spatial WM have shown that only a prioritized WM representation can be easily decoded in visual cortex, while a still-remembered but non-prioritized representation is indistinguishable from chance (Christophel et al., 2018; LaRocque et al., 2017, 2013; Lewis-Peacock et al., 2012; Lorenc et al., 2020; Rose et al., 2016; Sahan et al., 2019; Yu et al., 2020). These results have been interpreted as evidence that only items in the “focus of attention” (Cowan, 2001) can be decoded using multivariate methods.

If fMRI signals measured here and elsewhere are only sensitive to rehearsal via covert spatial attention, then we would not expect there to be a substantial impact of a task-irrelevant interrupting stimulus. Our results are consistent with this possibility. However, it is important to point out a wrinkle in this interpretation: a recent study which required participants to transiently attend to a cued location on the screen during spatial WM maintenance showed that, following a transient disruption of spatial WM representations in visual and parietal cortex, WM representations recovered substantially prior to the behavioral response. Interestingly, the authors correlated trial-by-trial errors in the decoded position and errors in each trial’s behavioral response and found a significant relationship only in retinotopic visual cortex (V1-V3), with no correlation observed in parietal or frontal cortex (Hallenbeck et al., 2021). This suggests a key role of local activation patterns in these regions in constraining behavioral performance on WM tasks. That is, if decoded WM representations in visual cortex merely reflect the impact of control or rehearsal processes occurring elsewhere, it is challenging to explain why only the target of the control processes in visual cortex exhibits a behaviorally-correlated bias, while the putative source of the control signals in parietal and frontal cortex does not exhibit these effects.

## Conclusions

We sought to characterize whether a brief, task-irrelevant interrupting stimulus could disrupt or augment spatial WM representations in retinotopic human cortex. While the interrupting stimulus was effective at evoking strong fMRI responses in all regions examined (**Fig. 2**), it did not result in a change in the fidelity of WM representations recovered using multiple implementations of an inverted encoding model (**Figs. 3-5**). The stability of these WM representations may be due to their redundancy across cortex, the particular integrity of spatial WM representations, or may suggest that fMRI signals are primarily sensitive to rehearsal or control processes. Altogether, these results place important constraints on the neural mechanisms supporting spatial WM, which can be further tested in future studies.

## Acknowledgments

We thank John Serences for providing funding to acquire data and for useful discussions at an early stage of this research.

## References

Awh E, Jonides J (2001) Overlapping mechanisms of attention and spatial working memory. Trends in Cognitive Sciences 5:119–126.

Awh E, Jonides J, Reuter-Lorenz PA (1998) Rehearsal in spatial working memory. Journal of Experimental Psychology: Human Perception and Performance 24:780.

Barbosa J, Soldevilla DL, Compte A (2021) Pinging reveals active, not silent, working memories.

Benjamini Y, Yekutieli D (2001) The control of the false discovery rate in multiple testing under dependency. Annals of statistics 29:1165–1188.

Bettencourt KC, Xu Y (2016) Decoding the content of visual short-term memory under distraction in occipital and parietal areas. Nat Neurosci 19:150–157.

Brainard DH (1997) The Psychophysics Toolbox. Spatial Vision 10:433–436.

Chafee MV, Goldman-Rakic PS (1998) Matching patterns of activity in primate prefrontal area 8a and parietal area 7ip neurons during a spatial working memory task. J Neurophysiol 79:2919–2940.

Chota S, Van der Stigchel S (2021) Dynamic and flexible transformation and reallocation of visual working memory representations. Visual Cognition 29:409–415.

Christophel TB, Cichy RM, Hebart MN, Haynes J-D (2015) Parietal and early visual cortices encode working memory content across mental transformations. Neuroimage 106:198–206.

Christophel TB, Haynes J-D (2014) Decoding complex flow-field patterns in visual working memory. NeuroImage 91C:43–51.

Christophel TB, Hebart MN, Haynes J-DJ-D (2012) Decoding the Contents of Visual Short-Term Memory from Human Visual and Parietal Cortex. The Journal of Neuroscience 32:12983–12989.

Christophel TB, Iamshchinina P, Yan C, Allefeld C, Haynes J-D (2018) Cortical specialization for attended versus unattended working memory. Nature Neuroscience 21:494–496.

Christophel TB, Klink PC, Spitzer B, Roelfsema PR, Haynes J-D (2017) The Distributed Nature of Working Memory. Trends Cogn Sci 21:111–124.

Courtney SM, Ungerleider LG, Keil K, Haxby JV (1997) Transient and sustained activity in a distributed neural system for human working memory. Nature 386:608–611.

Cowan N (2001) The magical number 4 in short-term memory: A reconsideration of mental storage capacity. Behavioral and Brain Sciences 24:87–114.

Curtis CE, D’Esposito M (2003) Persistent activity in the prefrontal cortex during working memory. Trends in cognitive sciences 7:415–423.

Curtis CE, Sprague TC (2021) Persistent Activity During Working Memory From Front to Back. Front Neural Circuits 15:696060.

Dubois J, de Berker AO, Tsao DY (2015) Single-unit recordings in the macaque face patch system reveal limitations of fMRI MVPA. The Journal of neuroscience : the official journal of the Society for Neuroscience 35:2791–802.

Emrich SM, Riggall AC, Larocque JJ, Postle BR (2013) Distributed patterns of activity in sensory cortex reflect the precision of multiple items maintained in visual short-term memory. The Journal of Neuroscience 33:6516–23.

Engel SA, Rumelhart DE, Wandell BA, Lee AT, Glover GH, Chichilnisky E-J, Shadlen MN (1994) fMRI of human visual cortex. Nature 369:525.

Ester EF, Anderson DE, Serences JT, Awh E (2013) A Neural Measure of Precision in Visual Working Memory. Journal of Cognitive Neuroscience 754–761.

Ester EF, Nouri A, Rodriguez L (2018) Retrospective cues mitigate information loss in human cortex during working memory storage. The Journal of neuroscience : the official journal of the Society for Neuroscience 1566–18.

Ester EF, Serences JT, Awh E (2009) Spatially Global Representations in Human Primary Visual Cortex during Working Memory Maintenance. The Journal of Neuroscience 29:15258–15265.

Ester EF, Sprague TC, Serences JT (2015) Parietal and Frontal Cortex Encode Stimulus-Specific Mnemonic Representations during Visual Working Memory. Neuron 87:893–905.

Foster JJ, Bsales EM, Jaffe RJ, Awh E (2017) Alpha-Band Activity Reveals Spontaneous Representations of Spatial Position in Visual Working Memory. Current Biology 27:3216-3223.e6.

Funahashi S, Bruce CJ, Goldman-Rakic PS (1989) Mnemonic coding of visual space in the monkey’s dorsolateral prefrontal cortex. Journal of neurophysiology 61:331–49.

Fuster JM, Alexander GE (1971) Neuron activity related to short-term memory. Science (New York, NY) 173:652–4.

Gayet S, Paffen CLE, Van der Stigchel S (2018) Visual Working Memory Storage Recruits Sensory Processing Areas. Trends in Cognitive Sciences 22:189–190.

Gnadt JW, Andersen RA (1988) Memory related motor planning activity in posterior parietal cortex of macaque. Exp Brain Res 70:216–220.

Hagler DJ, Sereno MI (2006) Spatial maps in frontal and prefrontal cortex. NeuroImage 29:567–77.

Hakim N, Feldmann-Wüstefeld T, Awh E, Vogel EK (2020) Perturbing Neural Representations of Working Memory with Task-irrelevant Interruption. J Cogn Neurosci 32:558–569.

Hallenbeck GE, Sprague TC, Rahmati M, Sreenivasan KK, Curtis CE (2021) Working memory representations in visual cortex mediate distraction effects. Nat Commun 12:4714.

Harrison SA, Tong F (2009) Decoding reveals the contents of visual working memory in early visual areas. Nature 458:632–635.

Henderson MM, Rademaker RL, Serences JT (2021) Flexible utilization of spatial-and motor-based codes for the storage of visuo-spatial information. bioRxiv 2021.07.08.451663.

Hermes D, Nguyen M, Winawer J (2017) Neuronal synchrony and the relation between the blood-oxygen-level dependent response and the local field potential. PLOS Biology 15:e2001461.

Iamshchinina P, Christophel TB, Gayet S, Rademaker RL (2021) Essential considerations for exploring visual working memory storage in the human brain. null 1–12.

Itthipuripat S, Sprague TC, Serences JT (2019) Functional MRI and EEG Index Complementary Attentional Modulations. J Neurosci 39:6162–6179.

Jerde TA, Merriam EP, Riggall AC, Hedges JH, Curtis CE (2012) Prioritized Maps of Space in Human Frontoparietal Cortex. The Journal of Neuroscience 32:17382–17390.

Kubota K, Niki H (1971) Prefrontal cortical unit activity and delayed alternation performance in monkeys. J Neurophysiol 34:337–347.

LaRocque JJ, Lewis-Peacock JA, Drysdale AT, Oberauer K, Postle BR (2013) Decoding attended information in short-term memory: an EEG study. Journal of cognitive neuroscience 25:127–42.

LaRocque JJ, Riggall AC, Emrich SM, Postle BR (2017) Within-Category Decoding of Information in Different Attentional States in Short-Term Memory. Cereb Cortex 27:4881–4890.

Leung H-C, Gore JC, Goldman-Rakic PS (2002) Sustained mnemonic response in the human middle frontal gyrus during on-line storage of spatial memoranda. J Cogn Neurosci 14:659–671.

Lewis-Peacock JA, Drysdale AT, Oberauer K, Postle BR (2012) Neural evidence for a distinction between short-term memory and the focus of attention. Journal of cognitive neuroscience 24:61–79.

Li H-H, Sprague TC, Yoo AH, Ma WJ, Curtis CE (2021) Joint representation of working memory and uncertainty in human cortex. Neuron 0.

Lorenc ES, Mallett R, Lewis-Peacock JA (2021) Distraction in Visual Working Memory: Resistance is Not Futile. Trends in Cognitive Sciences.

Lorenc ES, Sreenivasan KK (2021) Reframing the debate: The distributed systems view of working memory. Visual Cognition 0:1–9.

Lorenc ES, Sreenivasan KK, Nee DE, Vandenbroucke ARE, D’Esposito M (2018) Flexible Coding of Visual Working Memory Representations during Distraction. J Neurosci 38:5267–5276.

Lorenc ES, Vandenbroucke ARE, Nee DE, Lange FP de, D’Esposito M (2020) Dissociable neural mechanisms underlie currently-relevant, future-relevant, and discarded working memory representations. Sci Rep 10:1–17.

Mackey WE, Winawer J, Curtis CE (2017) Visual field map clusters in human frontoparietal cortex. eLife 6.

Macoveanu J, Klingberg T, Tegnér J (2007) Neuronal firing rates account for distractor effects on mnemonic accuracy in a visuo-spatial working memory task. Biol Cybern 96:407–419.

Mongillo G, Barak O, Tsodyks M (2008) Synaptic theory of working memory. Science (New York, NY) 319:1543–6.

Parthasarathy A, Herikstad R, Bong JH, Medina FS, Libedinsky C, Yen S-C (2017) Mixed selectivity morphs population codes in prefrontal cortex. Nat Neurosci 20:1770–1779.

Pasternak T, Greenlee MW (2005) Working memory in primate sensory systems. Nature Reviews Neuroscience 6:97–107.

Pelli DG (1997) The VideoToolbox software for visual psychophysics: transforming numbers into movies. Spatial Vision 10:437–442.

Postle BR, Yu Q (2020) Neuroimaging and the localization of function in visual cognition. Visual Cognition 0:1–6.

Rademaker RL, Bloem IM, De Weerd P, Sack AT (2015) The impact of interference on short-term memory for visual orientation. J Exp Psychol Hum Percept Perform 41:1650–1665.

Rademaker RL, Chunharas C, Serences JT (2019) Coexisting representations of sensory and mnemonic information in human visual cortex. Nature Neuroscience 22:1336–1344.

Riggall AC, Postle BR (2012) The relationship between working memory storage and elevated activity as measured with functional magnetic resonance imaging. The Journal of Neuroscience 32:12990–8.

Rose NS, LaRocque JJ, Riggall AC, Gosseries O, Starrett MJ, Meyering EE, Postle BR (2016) Reactivation of latent working memories with transcranial magnetic stimulation. Science 354:1136–1139.

Saber GT, Pestilli F, Curtis CE (2015) Saccade planning evokes topographically specific activity in the dorsal and ventral streams. The Journal of neuroscience : the official journal of the Society for Neuroscience 35:245–52.

Sahan MI, Sheldon AD, Postle BR (2019) The Neural Consequences of Attentional Prioritization of Internal Representations in Visual Working Memory. Journal of Cognitive Neuroscience 32:917–944.

Schneegans S, Bays PM (2017) Restoration of fMRI Decodability Does Not Imply Latent Working Memory States. J Cogn Neurosci 29:1977–1994.

Scimeca JM, Kiyonaga A, D’Esposito M (2018) Reaffirming the Sensory Recruitment Account of Working Memory. Trends in Cognitive Sciences 22:190–192.

Serences JT (2016) Neural mechanisms of information storage in visual short-term memory. Vision Research 128:53–67.

Serences JT, Ester EF, Vogel EK, Awh E (2009) Stimulus-Specific Delay Activity in Human Primary Visual Cortex. Psychological Science 20:207–214.

Sprague TC, Adam KCS, Foster JJ, Rahmati M, Sutterer DW, Vo VA (2018a) Inverted Encoding Models Assay Population-Level Stimulus Representations, Not Single-Unit Neural Tuning. eNeuro 5.

Sprague TC, Boynton GM, Serences JT (2019) The Importance of Considering Model Choices When Interpreting Results in Computational Neuroimaging. eNeuro 6.

Sprague TC, Ester EF, Serences JT (2016) Restoring Latent Visual Working Memory Representations in Human Cortex. Neuron 91:694–707.

Sprague TC, Ester EF, Serences JT (2014) Reconstructions of Information in Visual Spatial Working Memory Degrade with Memory Load. Current Biology.

Sprague TC, Itthipuripat S, Vo VA, Serences JT (2018b) Dissociable signatures of visual salience and behavioral relevance across attentional priority maps in human cortex. Journal of Neurophysiology 119:2153–2165.

Sprague TC, Serences JT (2013) Attention modulates spatial priority maps in the human occipital, parietal and frontal cortices. Nature neuroscience 16:1879–87.

Sreenivasan KK, D’Esposito M (2019) The what, where and how of delay activity. Nature Reviews Neuroscience 20:466–481.

Srimal R, Curtis CE (2008) Persistent neural activity during the maintenance of spatial position in working memory. NeuroImage 39:455–468.

Stokes MG (2015) “Activity-silent” working memory in prefrontal cortex: a dynamic coding framework. Trends in cognitive sciences 19:394–405.

Supèr H, Spekreijse H, Lamme VA (2001) A neural correlate of working memory in the monkey primary visual cortex. Science 293:120–124.

Swisher JD, Halko MA, Merabet LB, McMains SA, Somers DC (2007) Visual topography of human intraparietal sulcus. The Journal of Neuroscience 27:5326–5337.

Teng C, Postle BR (2021) Understanding occipital and parietal contributions to visual working memory: Commentary on Xu (2020). Visual Cognition 0:1–8.

van Kerkoerle T, Self MW, Roelfsema PR (2017) Layer-specificity in the effects of attention and working memory on activity in primary visual cortex. Nature Communications 8:13804.

van Moorselaar D, Foster JJ, Sutterer DW, Theeuwes J, Olivers CNL, Awh E (2018) Spatially Selective Alpha Oscillations Reveal Moment-by-Moment Trade-offs between Working Memory and Attention. Journal of Cognitive Neuroscience 30:256–266.

Wolff MJ, Akyurek E, Stokes MG (2021) What is the functional role of delay-related alpha oscillations during working memory?

Wolff MJ, Ding J, Myers NE, Stokes MG (2015) Revealing hidden states in visual working memory using electroencephalography. Frontiers in systems neuroscience 9:123.

Wolff MJ, Jochim J, Akyürek EG, Buschman TJ, Stokes MG (2020) Drifting codes within a stable coding scheme for working memory. PLOS Biology 18:e3000625.

Wolff MJ, Jochim J, Akyürek EG, Stokes MG (2017) Dynamic hidden states underlying working-memory-guided behavior. Nature Neuroscience 20:864–871.

Xu Y (2021) Towards a better understanding of information storage in visual working memory. Visual Cognition 0:1–9.

Xu Y (2020) Revisit once more the sensory storage account of visual working memory. Visual Cognition 0:1–14.

Xu Y (2018) Sensory Cortex Is Nonessential in Working Memory Storage. Trends in Cognitive Sciences 22:192–193.

Xu Y (2017) Reevaluating the Sensory Account of Visual Working Memory Storage. Trends in Cognitive Sciences 21:794–815.

Yu Q, Shim WM (2017) Occipital, parietal, and frontal cortices selectively maintain task-relevant features of multi-feature objects in visual working memory. NeuroImage 157:97–107.

Yu Q, Teng C, Postle BR (2020) Different states of priority recruit different neural representations in visual working memory. PLOS Biology 18:e3000769.

